# An improved mitochondrial reference genome for *Arabidopsis thaliana Col-0*

**DOI:** 10.1101/249086

**Authors:** Daniel B. Sloan, Zhiqiang Wu, Joel Sharbrough

**Affiliations:** Department of Biology, Colorado State University, Fort Collins, CO 80523

## Abstract

*Arabidopsis thaliana* remains the foremost model system for plant genetics and genomics, and researchers rely on the accuracy of its genomic resources. The first completely sequenced angiosperm mitochondrial genome was obtained from *A. thaliana C24* (Unseld et al., 1997), and more recent efforts have produced additional *A. thaliana* reference genomes, including one for *Col-0*, the most widely used ecotype (Davila et al., 2011). These studies were based on older DNA sequencing methods, making them subject to errors associated with lower levels of sequencing coverage or the extremely short read lengths produced by early-generation Illumina technologies. Indeed, although the more recently published *A. thaliana* mitochondrial reference genome sequences made substantial progress in improving upon earlier versions, they still have high error rates. By comparing publicly available Illumina sequence data to the *A. thaliana Col-0* reference genome, we found that it contains a sequence error every 2.4 kb on average, including 57 SNPs, 96 indels (up to 901 bp in size), and a large repeat-mediated rearrangement. Most of these errors appear to have been carried over from the original *A. thaliana* mitochondrial genome sequence by reference-based assembly approaches, which has misled subsequent studies of plant mitochondrial mutation and molecular evolution by giving the false impression that the errors are naturally occurring variants present in multiple ecotypes. Building on the progress made by previous researchers, we provide a corrected reference sequence that we hope will serve as a useful community resource for future investigations in the field of plant mitochondrial genetics.

## The history of *Arabidopsis* mitochondrial genome sequencing

In 1997, a group led by Axel Brennicke reported the landmark achievement of sequencing a complete mitochondrial genome from *A. thaliana* (Unseld et al., 1997), ushering flowering plants into the era of mitogenomics and providing numerous insights about the distinctive features of mitochondrial DNA (mtDNA) in plants (Mower et al., 2012). There has been some confusion over the source material used for this first sequencing effort. In the original 1997 publication and current GenBank accessions (Y08501.2 and NC_001284.2), the source ecotype is described as *Columbia*. However, the cosmid library used for sequencing was derived from *C24* (Klein et al., 1994), which is genetically distinct from the widely used *Col-0* (i.e., *Columbia*) ecotype (Lehle Seeds, 2004). The *C24* source of the original published genome has been confirmed in subsequent studies (Davila et al., 2011). Nevertheless, some confusion persists in research that has misinterpreted the *C24* sequence as being from *Col-0* (e.g., Zampini et al., 2015).

More recent efforts in the early phases of the “next-generation” sequencing revolution resequenced the mitochondrial genome of the *C24* ecotype (GenBank accession JF729200) and produced reference sequences for the *Col-0* (JF729201) and *Ler* (JF729202) ecotypes (Davila et al., 2011). Resequencing of *C24* yielded the same overall genome structure as the original sequence (Unseld et al., 1997) and earlier mapping efforts (Klein et al., 1994), but it also produced 416 sequence differences in the form of SNPs and small indels. At the time, there was no discussion or further investigation of these sequence differences, but they appear to represent corrections of sequencing errors from the original genome rather than true biological differences. Therefore, the work by Davila et al. (2011) has led to valuable increases and improvements in available mitogenomic resources for *A. thaliana*. However, these efforts relied on some of the earliest implementations of Illumina sequencing technology. The extremely short read-lengths (35 bp) that were available at the onset of that study limited the researchers to reference-based assembly approaches, posing challenges for identification of variants in regions with multiple sequence differences. Therefore, the accuracy of the available *A. thaliana* reference genomes has remained uncertain.

## Persistent sequencing errors in published *Arabidopsis* mitochondrial genomes

While performing research to identify naturally occurring variants in *A. thaliana* mtDNA (and being ignorant of some of the history described above), we were surprised to find that sequence datasets from *A. thaliana Col-0* exhibited numerous mitochondrial variants even when mapped against the *Col-0* reference sequence. To investigate these discrepancies, we used a publicly available Illumina MiSeq dataset (2 × 300-bp paired-end reads; NCBI SRA SRR5216995) to perform a *de novo* assembly of the *A. thaliana Col-0* mitochondrial genome by employing the SPAdes Genome Assembler v3.11.0 (Bankevich et al., 2012) with a range of ft-mers (21, 33, 55, 77, and 99) followed by manual inspection and joining of contigs. The resulting assembly differed by 57 SNPs and 96 indels relative to the published *A. thaliana Col-0* reference genome (Davila et al., 2011), amounting to a variant every 2.4 kb on average. To assess whether these variants represented sequencing artefacts or actual biological differences between the two *Col-0* samples, we extracted diagnostic *k*-mers from the raw reads used in our analysis and those from the original *A. thaliana Col-0* sequencing effort (SRA SRR307226). We confirmed that all the variants identified in our assembly were strongly supported in both sets of sequencing reads (Table 1), suggesting that the differences represent assembly errors in the published *Col-0* reference sequence rather than real polymorphisms. We further validated these variants calls using the double-stranded consensus sequence from a dataset (SRA SRR6420475) that was generated with a highly accurate technique known as duplex sequencing (Schmitt et al., 2012).

**Table 1.** *k*-mer based support for corrected *A. thaliana Col-0* sequence. Illumina read counts are based on presence of diagnostic *k*-mers that distinguish between the reported variants in *Col-0* and *C24* sequence datasets. The reference genome and corrected *Col-0* genomes correspond to accessions JF729201.1 and BK010421, respectively. SRR5216995 is the raw dataset used for our *de novo* assembly. SRR307226 and SRR307231 are the raw datasets produced by Davila et al. (2011) for *Col-0* and *C24*, respectively.

| Ref Pos | Corr Pos | Type | Length | Ref Allele | Corr Allele | SRR5216995 Read Counts |  | SRR307226 Read Counts |  | SRR307231 Read Counts |  |
| --- | --- | --- | --- | --- | --- | --- | --- | --- | --- | --- | --- |
|  |  |  |  |  |  | Ref | Corrected | Ref | Corrected | Ref | Corrected |
| 844 | 845 | Ins | 1 |  | C | 0 | 655 | 0 | 189 | 0 | 321 |
| 1179 | 1179 | Del | 5 | TGTTT |  | 0 | 743 | 1 | 131 | 203 | 17 |
| 13043 | 13040 | Ins | 5 |  | AGAAT | 0 | 514 | 0 | 55 | 117 | 9 |
| 19275 | 19277 | Ins | 1 |  | G | 0 | 79 | 0 | 40 | 0 | 48 |
| 19288 | 19290 | SNP | 1 | T | G | 0 | 57 | 0 | 26 | 25 | 5 |
| 21599 | 21602 | Ins | 1 |  | T | 0 | 345 | 0 | 62 | 0 | 105 |
| 26459 | 26463 | Ins | 1 |  | G | 0 | 461 | 0 | 39 | 0 | 82 |
| 27254 | 27259 | Ins | 1 |  | G | 0 | 368 | 0 | 73 | 0 | 116 |
| 27983 | 27989 | Ins | 1 |  | A | 0 | 394 | 0 | 86 | 0 | 130 |
| 28469 | 28476 | Ins | 1 |  | G | 0 | 511 | 0 | 60 | 0 | 119 |
| 32254 | 32262 | Ins | 1 |  | G | 0 | 539 | 0 | 66 | 0 | 124 |
| 38425 | 142337 | Ins | 1 |  | C | 0 | 429 | 0 | 104 | 0 | 157 |
| 40270 | 140491 | Ins | 1 |  | C | 0 | 547 | 1 | 85 | 0 | 102 |
| 48894 | 131866 | Ins | 901 |  | 901 bp | 0 | 628 | 0 | 74 | 0 | 119 |
| 51009 | 128850 | Ins | 1 |  | T | 9 | 322 | 2 | 60 | 1 | 106 |
| 52038 | 127820 | Ins | 1 |  | C | 0 | 309 | 0 | 70 | 0 | 120 |
| 54300 | 125557 | Ins | 1 |  | T | 0 | 544 | 0 | 89 | 0 | 143 |
| 58720 | 121136 | Ins | 9 |  | 9 bp | 0 | 432 | 0 | 99 | 0 | 152 |
| 69330 | 110517 | Ins | 1 |  | T | 0 | 794 | 0 | 174 | 0 | 258 |
| 69720 | 110128 | Del | 1 | T |  | 0 | 308 | 1 | 111 | 5 | 146 |
| 81344 | 98505 | Del | 1 | G |  | 0 | 457 | 0 | 84 | 0 | 110 |
| 81755 | 98095 | Del | 1 | G |  | 0 | 407 | 0 | 80 | 0 | 116 |
| 82131 | 97718 | Ins | 1 |  | A | 0 | 336 | 0 | 97 | 1 | 139 |
| 82653 | 97195 | Ins | 1 |  | T | 0 | 568 | 0 | 76 | 0 | 112 |
| 83180 | 96668 | SNP | 1 | T | A | 0 | 544 | 0 | 60 | 0 | 111 |
| 83181 | 96667 | SNP | 1 | T | A | 0 | 546 | 0 | 58 | 0 | 119 |
| 83183 | 96665 | SNP | 1 | A | T | 0 | 552 | 0 | 61 | 0 | 105 |
| 83184 | 96664 | SNP | 1 | A | T | 0 | 551 | 0 | 63 | 0 | 108 |
| 85683 | 94164 | Ins | 4 |  | GCGC | 0 | 425 | 0 | 56 | 0 | 92 |
| 107160 | 72683 | Ins | 1 |  | T | 0 | 535 | 0 | 89 | 0 | 131 |
| 107614 | 72228 | Ins | 1 |  | T | 0 | 616 | 0 | 87 | 0 | 141 |
| 112983 | 66858 | Ins | 31 |  | 31 bp | 0 | 587 | 0 | 79 | 0 | 86 |
| 117081 | 62729 | Ins | 2 |  | GC | 0 | 467 | 0 | 98 | 0 | 146 |
| 122998 | 56812 | Del | 1 | C |  | 0 | 411 | 0 | 61 | 0 | 109 |
| 130836 | 48973 | Ins | 1 |  | G | 0 | 792 | 0 | 168 | 0 | 243 |
| 131249 | 48559 | Ins | 1 |  | T | 0 | 448 | 0 | 68 | 0 | 100 |
| 131787 | 48020 | Ins | 1 |  | G | 0 | 440 | 0 | 93 | 0 | 132 |
| 131824 | 47982 | Ins | 1 |  | A | 0 | 454 | 0 | 83 | 0 | 135 |
| 133190 | 46617 | Del | 1 | T |  | 0 | 573 | 0 | 132 | 0 | 192 |
| 135089 | 44718 | SNP | 1 | C | T | 0 | 1860 | 0 | 236 | 0 | 321 |
| 135125 | 44682 | SNP | 1 | G | C | 2 | 1430 | 0 | 179 | 0 | 194 |
| 135167 | 44639 | Ins | 2 |  | TC | 0 | 482 | 0 | 56 | 0 | 95 |
| 135631 | 44173 | Ins | 1 |  | G | 0 | 388 | 0 | 93 | 0 | 112 |
| 136553 | 43250 | Ins | 6 |  | TAAATT | 0 | 552 | 0 | 50 | 47 | 6 |
| 137330 | 42467 | Ins | 1 |  | A | 0 | 363 | 0 | 74 | 0 | 97 |
| 139304 | 40492 | Ins | 1 |  | G | 0 | 376 | 0 | 84 | 0 | 189 |
| 142386 | 37409 | Ins | 1 |  | A | 0 | 500 | 0 | 100 | 0 | 165 |
| 143342 | 36452 | Ins | 1 |  | G | 0 | 918 | 0 | 157 | 0 | 232 |
| 146050 | 147026 | Del | 3 | TTA |  | 0 | 393 | 0 | 72 | 0 | 165 |
| 146054 | 147028 | SNP | 1 | G | A | 0 | 390 | 0 | 74 | 0 | 169 |
| 146055 | 147029 | SNP | 1 | G | T | 0 | 389 | 0 | 79 | 0 | 167 |
| 146057 | 147031 | SNP | 1 | A | G | 0 | 384 | 0 | 99 | 0 | 172 |
| 146058 | 147032 | SNP | 1 | T | A | 0 | 381 | 0 | 103 | 0 | 168 |
| 146073 | 147046 | Del | 1 | T |  | 0 | 376 | 0 | 83 | 0 | 162 |
| 146080 | 147052 | Del | 5 | TATTG |  | 0 | 371 | 0 | 84 | 0 | 167 |
| 146087 | 147055 | SNP | 1 | T | G | 0 | 365 | 0 | 95 | 0 | 182 |
| 146089 | 147057 | SNP | 1 | C | T | 0 | 360 | 0 | 86 | 0 | 197 |
| 146090 | 147058 | SNP | 1 | C | A | 0 | 364 | 0 | 94 | 0 | 203 |
| 146091 | 147059 | SNP | 1 | G | C | 0 | 366 | 0 | 94 | 0 | 205 |
| 146094 | 147062 | SNP | 1 | A | G | 0 | 366 | 0 | 93 | 0 | 202 |
| 146100 | 147067 | Del | 3 | TAA |  | 0 | 369 | 0 | 84 | 0 | 175 |
| 154362 | 155328 | Ins | 1 |  | A | 0 | 466 | 1 | 73 | 1 | 117 |
| 156407 | 157374 | Ins | 1 |  | C | 0 | 506 | 0 | 99 | 0 | 136 |
| 157813 | 158779 | Del | 1 | G |  | 0 | 480 | 0 | 101 | 0 | 139 |
| 163337 | 164304 | Ins | 1 |  | A | 0 | 487 | 0 | 66 | 0 | 129 |
| 164285 | 165251 | Del | 1 | A |  | 0 | 6457 | 0 | 192 | 0 | 325 |
| 173702 | 174667 | Del | 1 | G |  | 0 | 428 | 0 | 56 | 0 | 51 |
| 178699 | 179665 | Ins | 1 |  | T | 0 | 489 | 0 | 75 | 0 | 94 |
| 194075 | 195042 | Ins | 1 |  | C | 0 | 655 | 0 | 189 | 0 | 321 |
| 194410 | 195376 | Del | 5 | TGTTT |  | 0 | 743 | 1 | 131 | 203 | 17 |
| 220011 | 220974 | Ins | 99 |  | 99 bp | 0 | 445 | 0 | 75 | 0 | 122 |
| 221620 | 222680 | Del | 4 | TCCT |  | 0 | 895 | 0 | 169 | 0 | 113 |
| 222813 | 223871 | Ins | 1 |  | C | 4 | 43 | 6 | 19 | 41 | 5 |
| 226207 | 227266 | Ins | 1 |  | C | 0 | 425 | 0 | 63 | 0 | 92 |
| 232692 | 233750 | Del | 1 | A |  | 0 | 343 | 0 | 76 | 0 | 121 |
| 233251 | 234309 | SNP | 1 | C | A | 0 | 857 | 0 | 88 | 0 | 66 |
| 233258 | 234315 | Del | 1 | T |  | 0 | 853 | 0 | 87 | 0 | 71 |
| 235638 | 236696 | Ins | 50 |  | 50 bp | 0 | 291 | 0 | 69 | 0 | 104 |
| 235958 | 237066 | Ins | 1 |  | C | 7 | 57 | 6 | 25 | 36 | 6 |
| 238603 | 239710 | Del | 1 | A |  | 0 | 405 | 0 | 67 | 0 | 114 |
| 238663 | 239771 | Ins | 1 |  | G | 0 | 431 | 0 | 57 | 0 | 95 |
| 239186 | 240295 | Ins | 1 |  | T | 0 | 348 | 0 | 78 | 0 | 143 |
| 254302 | 255412 | Ins | 1 |  | A | 0 | 416 | 0 | 79 | 0 | 137 |
| 254353 | 255464 | Ins | 1 |  | C | 0 | 392 | 0 | 42 | 0 | 94 |
| 254424 | 255535 | SNP | 1 | G | T | 0 | 415 | 0 | 84 | 0 | 110 |
| 254430 | 255541 | SNP | 1 | A | G | 0 | 408 | 0 | 69 | 0 | 110 |
| 254431 | 255543 | Ins | 1 |  | C | 0 | 411 | 0 | 70 | 0 | 110 |
| 254473 | 255585 | SNP | 1 | G | A | 0 | 445 | 0 | 61 | 0 | 122 |
| 254474 | 255586 | SNP | 1 | A | G | 0 | 447 | 0 | 59 | 0 | 122 |
| 254475 | 255588 | Ins | 2 |  | AT | 0 | 448 | 0 | 54 | 0 | 116 |
| 254484 | 255598 | SNP | 1 | C | T | 0 | 427 | 0 | 61 | 0 | 126 |
| 254485 | 255599 | SNP | 1 | G | C | 0 | 424 | 0 | 60 | 0 | 128 |
| 254489 | 255603 | SNP | 1 | A | G | 0 | 423 | 0 | 54 | 0 | 137 |
| 254489 | 255604 | Ins | 4 |  | AATC | 0 | 423 | 0 | 54 | 0 | 136 |
| 254491 | 255609 | SNP | 1 | C | T | 0 | 425 | 0 | 56 | 0 | 132 |
| 254492 | 255610 | SNP | 1 | T | G | 0 | 426 | 0 | 59 | 0 | 133 |
| 254493 | 255611 | SNP | 1 | G | C | 0 | 423 | 0 | 59 | 0 | 130 |
| 254494 | 255612 | SNP | 1 | C | A | 0 | 422 | 0 | 61 | 0 | 133 |
| 254504 | 255623 | Ins | 1 |  | A | 0 | 431 | 0 | 59 | 0 | 119 |
| 254918 | 256036 | Del | 48 | 48 bp |  | 0 | 415 | 0 | 16 | 17 | 1 |
| 256144 | 257214 | Del | 1 | C |  | 0 | 426 | 0 | 66 | 0 | 84 |
| 264495 | 265565 | SNP | 1 | C | G | 0 | 1291 | 0 | 205 | 91 | 264 |
| 265386 | 266457 | Ins | 1 |  | C | 0 | 478 | 0 | 64 | 0 | 119 |
| 266189 | 267261 | Ins | 1 |  | A | 0 | 487 | 0 | 88 | 169 | 7 |
| 267980 | 269053 | Ins | 1 |  | T | 0 | 415 | 0 | 64 | 0 | 104 |
| 268497 | 269569 | Del | 32 | 32 bp |  | 0 | 466 | 0 | 69 | 0 | 109 |
| 276590 | 277632 | Ins | 3 |  | TGC | 0 | 379 | 0 | 65 | 0 | 12 |
| 282381 | 283425 | SNP | 1 | G | A | 0 | 573 | 0 | 86 | 0 | 125 |
| 282381 | 283426 | Ins | 2 |  | GA | 0 | 573 | 0 | 88 | 0 | 127 |
| 282383 | 283429 | SNP | 1 | A | G | 0 | 572 | 0 | 87 | 0 | 129 |
| 282384 | 283430 | SNP | 1 | G | A | 0 | 577 | 0 | 89 | 0 | 128 |
| 289149 | 290196 | Ins | 1 |  | T | 0 | 598 | 0 | 27 | 0 | 32 |
| 289699 | 290747 | Ins | 1 |  | T | 0 | 587 | 0 | 73 | 0 | 90 |
| 289713 | 290762 | Ins | 1 |  | A | 0 | 579 | 0 | 79 | 0 | 102 |
| 306032 | 307081 | SNP | 1 | G | A | 0 | 965 | 1 | 110 | 79 | 82 |
| 306058 | 307107 | SNP | 1 | T | C | 0 | 964 | 0 | 150 | 85 | 126 |
| 306061 | 307110 | SNP | 1 | A | G | 0 | 968 | 0 | 150 | 91 | 120 |
| 306062 | 307111 | SNP | 1 | G | T | 0 | 963 | 0 | 145 | 86 | 117 |
| 306063 | 307112 | SNP | 1 | C | T | 0 | 961 | 0 | 141 | 86 | 111 |
| 306685 | 307734 | SNP | 1 | A | G | 0 | 416 | 0 | 65 | 0 | 10 |
| 306689 | 307738 | SNP | 1 | A | G | 0 | 422 | 0 | 63 | 0 | 8 |
| 306692 | 307741 | SNP | 1 | A | G | 0 | 431 | 0 | 57 | 86 | 8 |
| 306693 | 307742 | SNP | 1 | A | G | 0 | 435 | 0 | 51 | 86 | 8 |
| 306694 | 307743 | SNP | 1 | C | G | 0 | 434 | 0 | 50 | 93 | 8 |
| 306701 | 307750 | SNP | 1 | A | G | 0 | 449 | 0 | 45 | 92 | 10 |
| 306704 | 307753 | SNP | 1 | A | C | 0 | 452 | 0 | 56 | 88 | 11 |
| 306707 | 307756 | SNP | 1 | A | G | 0 | 1145 | 0 | 124 | 88 | 116 |
| 306708 | 307757 | SNP | 1 | T | C | 0 | 1140 | 0 | 123 | 82 | 117 |
| 306709 | 307758 | SNP | 1 | T | A | 0 | 1137 | 0 | 123 | 75 | 115 |
| 306712 | 307761 | SNP | 1 | C | A | 0 | 1142 | 0 | 134 | 72 | 121 |
| 306713 | 307762 | SNP | 1 | G | A | 0 | 1135 | 0 | 131 | 80 | 119 |
| 306718 | 307767 | SNP | 1 | A | T | 0 | 1143 | 0 | 115 | 0 | 137 |
| 306720 | 307769 | SNP | 1 | C | T | 0 | 1148 | 0 | 130 | 0 | 140 |
| 306725 | 307774 | SNP | 1 | T | A | 0 | 1140 | 0 | 120 | 0 | 119 |
| 306726 | 307775 | SNP | 1 | T | C | 0 | 662 | 0 | 58 | 0 | 11 |
| 309015 | 310065 | Ins | 1 |  | G | 0 | 479 | 0 | 58 | 0 | 107 |
| 309250 | 310301 | Ins | 1 |  | G | 0 | 597 | 0 | 73 | 0 | 140 |
| 313181 | 314231 | Del | 2 | TC |  | 0 | 519 | 0 | 57 | 0 | 127 |
| 320937 | 321987 | Ins | 1 |  | G | 0 | 428 | 0 | 70 | 0 | 132 |
| 330316 | 331365 | Del | 1 | G |  | 0 | 410 | 0 | 48 | 0 | 10 |
| 331574 | 332623 | SNP | 1 | G | A | 24 | 314 | 0 | 63 | 0 | 12 |
| 332107 | 333157 | Ins | 1 |  | C | 0 | 500 | 0 | 55 | 0 | 11 |
| 339070 | 340121 | Ins | 1 |  | A | 2 | 393 | 0 | 66 | 0 | 98 |
| 339885 | 340937 | Ins | 1 |  | C | 2 | 352 | 0 | 54 | 0 | 98 |
| 340230 | 341283 | Ins | 1 |  | G | 0 | 415 | 0 | 71 | 0 | 127 |
| 343345 | 344399 | Ins | 1 |  | G | 0 | 464 | 0 | 84 | 0 | 95 |
| 347688 | 348743 | Ins | 1 |  | C | 1 | 425 | 0 | 67 | 0 | 99 |
| 354055 | 355111 | Ins | 1 |  | T | 0 | 472 | 0 | 73 | 0 | 137 |
| 358190 | 359246 | SNP | 1 | G | A | 4 | 381 | 1 | 42 | 62 | 5 |
| 358193 | 359249 | SNP | 1 | T | C | 4 | 381 | 1 | 42 | 62 | 5 |
| 358195 | 359251 | SNP | 1 | T | C | 4 | 381 | 1 | 42 | 62 | 5 |
| 358736 | 359793 | Ins | 1 |  | C | 0 | 266 | 0 | 52 | 0 | 75 |
| 361869 | 362927 | Ins | 1 |  | T | 0 | 350 | 0 | 61 | 0 | 110 |

By comparing the same set of 57 SNPs and 96 indels to the raw reads in the resequenced *C24* dataset (SRA SRR307231), we identified 28 variants for which the original reference allele was supported in *C24* (Table 1). These cases, therefore, represent true polymorphisms that distinguish the *C24* and *Col-0* ecotypes but were not detected in the original reference-based assembly of the *Col-0* mitochondrial genome such that the published *Col-0* sequence improperly retains the *C24* allele. In contrast, we found that the raw *C24* sequence reads did not support the original reference allele in the remaining 125 variants (82%) (Table 1). These cases appear to result from errors in the original *C24* genome sequence (Unseld et al., 1997) that were not detected in either the resequencing of *C24* or the reference-based assembly of *Col-0* and, thus, have been propagated across reported genome sequences from multiple ecotypes (Davila et al., 2011). Many of these errors are found in regions differing by multiple SNPs or by multi-nucleotide indels, so it is not surprising that they were difficult to detect with short-read sequencing data. However, there are also many individual SNPs and 1-bp indels in this set (Table 1), so the source of the assembly artefacts is unclear in some cases.

**Table 1.**
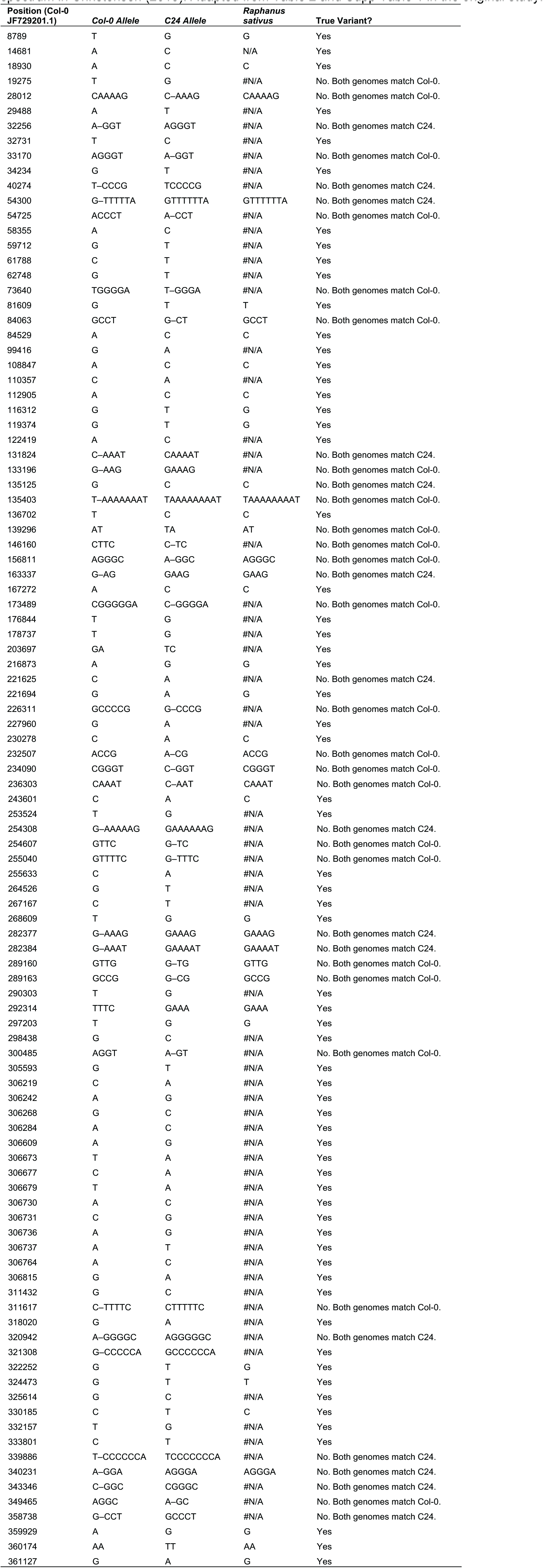
Identification of sequencing artefacts in comparative dataset used to infer mutational spectrum in Christensen (2013). Adapted from Table 2 and Supp Table 1 in the original study.

Our newly assembled *A. thaliana Col-0* reference sequence also differs from the published *Col-0* sequence in two major structural variants. First, it includes a 901-bp sequence that is absent from the published *Col-0* genome. The full-length of this sequence is clearly detectable in the raw reads of the original *Col-0* study (SRA SRR307226). It would be inserted after position 48,895 in the published *Col-0* genome (JF729201) and would correspond precisely to the last 901 bp of the *C24* reference genomes. The fact that this deletion occurs exactly at the point where the circular reference genome map had been arbitrarily “cut” for reporting as a linearized sequence suggests that it might have resulted from an inadvertent byproduct of sequence handling and reorientation. Second, our newly assembled *A. thaliana Col-0* reference sequence differs in a large rearrangement, apparently resulting from recombination between a pair of identical 453-bp inverted repeats at positions 36,362-36,818 and 143,953-144,409. The clear majority (30 of 33; 91%) of read-pairs spanning these repeats support our reported conformation. We are not able to test for similar support in the raw *Col-0* reads from Davila et al. (2011) because their insert sizes are too short to span the repeat copies, but we did verify that our reported configuration predominates in Illumina paired-end and PacBio sequencing reads from four other *Col-0* datasets (NCBI SRA SRR1581142, SRR5012968, SRR5882797, and SRP073602). Therefore, this configuration is likely the most common among different *Col-0* seed stocks.

## Subsequent research in *Arabidopsis* mitochondrial genetics

For good reason, *A. thaliana* is the “go-to” model for studies of plant mitochondrial genome function, stability, mutation, and molecular evolution (Davila et al., 2011; Christensen, 2013; Cupp and Nielsen, 2014; Zampini et al., 2015; Gualberto and Newton, 2017). As such, there is great incentive to make the *Arabidopsis* reference mitochondrial genomes the gold standard in the field. Indeed, the extensive characterization of structural variation in these genomes has gone a long way to accomplish this goal (Arrieta-Montiel et al., 2009). However, sequence errors still exist in the reported reference genomes with potentially detrimental and far-reaching effects on related research efforts. This is especially true because the actual rate of sequence evolution in plant mtDNA is usually very low (Wolfe et al., 1987), so even a modest amount of sequencing errors can result in a problematic signal-to-noise ratio. For example, a recent study was performed to infer the distribution and spectrum of mutations across the *Arabidopsis* mitochondrial genome and used the sequence variants that distinguish published *C24* and *Col-0* mtDNA sequences (Christensen, 2013). Such comparative analyses of published genomic data are commonplace and can make substantial contributions to the field, but it is now clear based on our reexamination of the *Col-0* sequence that approximately 40% of the analyzed variants in that study were artefacts (Table 2).

Another recent investigation was conducted to detect *de novo* mutations in *A. thaliana* organelle genomes using deep sequencing (Zampini et al., 2015). The authors applied a natural and seemingly conservative approach by rejecting any identified mitochondrial variant that did not differ from “both” published *Col-0* mitochondrial genomes, but this choice highlights two pressing concerns. First, it illustrates the continued confusion in the field about the fact that original *A. thaliana* reference mitochondrial genome is derived from *C24* and not *Col-0*. Second, it reflects a misunderstanding about the extent to which the multiple available reference genomes constitute independent data points. The reference-guided approach used to assemble mtDNA sequences from *C24, Col-0*, and *Ler* (Davila et al., 2011) appears to have incorporated many errors and allelic variants from the reference genome into the new assemblies. Nevertheless, those new assemblies are still reported as separate accessions on GenBank rather than as a set of variant calls, so there is a risk that the many errors shared between them will be falsely perceived as having been independently validated in two or more sequencing datasets. This concern is particularly relevant for the *Ler* sequence available on GenBank because it was generated with the same short 35-bp reads but a much lower level of sequence coverage – only 19× compared to 230× and 371 × for *Col-0* and *C24*, respectively (Davila et al., 2011). For these reasons, it is important that researchers in the field of plant mitochondrial genetics be more broadly aware of the history and methodologies that produced the currently available reference mitochondrial genome sequence for *A. thaliana*.

We have deposited our *de novo* assembly of the *A. thaliana Col-0* genome on GenBank (accession BK010421) in hopes that it will serve the community as a useful reference such that *A. thaliana* can further develop as an outstanding model for elucidating mitochondrial genetic mechanisms.

## Acknowledgements

This research was supported by the National Institutes of Health (NIGMS R01 GM118046).

## Author Contributions

D.B.S., Z.W., and J.S. performed data analysis. D.B.S. wrote the manuscript.

